# Influencing Iron Homeostasis by Transition Metal Complexes Interacting with a messenger-RNA Stem Loop in the Iron Regulating Element

**DOI:** 10.1101/2025.02.19.639007

**Authors:** Cécilia Hognon, Aurane Froux, Stéphanie Grandemange, Antonio Monari

## Abstract

The maintaining of the precise regulation of iron homeostasis is fundamental to assure cells viability. In this context the interplay between the messenger RNA Iron Response element (IRE) and the iron response protein 1 (IRP1) is crucial to regulate the expression of ferritin and hence the level of labile free iron pool. We have shown, using a combination of molecular modeling and experimental techniques, that tris bipyridine iron complexes (AIM3) interact specifically with the IRE RNA stem-loop in solution, without inducing noticeable structural deformations. Furthermore, we have also shown that, at a cellular level, this interaction may be traced back to the downregulation of ferritin translation, probably due to the stabilization of the IRE stem-loops thus favoring its binding to IRP1 or by inhibiting the downstream recruitment of ribosomal subunits.

## 1. Introduction

The precise maintaining of iron homeostasis is crucial to assure the cell survival and avoid the emergence of serious pathological conditions [1–3]. In cells, iron is usually stored in different proteins or nanocages, although a small amount of labile free iron pool (FIP) level is maintained and is required to assure cell viability. However, iron accumulation and the increase of FIP level may lead to serious outcomes, including oxidative stress [4–6]. In turn, these may ultimately result in increased cytotoxicity or mutagenesis [7,8] and be linked to neurodegenerative disorders [8–10] or cancer development [11–14]. The maintenance of FIP homeostasis is also complicated by the fact that iron concentration may vary depending on the environmental conditions experienced by the cells [15]. For instance, iron concentration may increase in presence of nutrients, and decrease in starving conditions [16]. Thus, cells should dispose of a regulatory mechanism, which is able to rapidly and promptly adapt responding to external stimuli. Iron homeostasis is controlled by a variety of different proteins, allowing iron internalization (transferrin receptor) [17,18], expulsion and outtake (ferroportin) [19,20], or storage (Ferritin H and L) [21,22]. Indeed, one of the most crucial endpoints of the iron regulation mechanisms involves the action of ferritin, a cytosolic globular protein allowing to diminish the free iron pool (FIP) content in cells by absorbing iron into nanocages presenting multiple metal/protein interaction sites [23,24]. Interestingly, the regulatory mechanisms take place at a translational level, and ferritin is overexpressed when the iron intracellular concentration is high, allowing the reduction of the FIP, while its translation is inhibited in case of iron depletion.

This regulatory mechanism (see Figure 1) is assured by the crosstalk between the Iron Response Element (IRE) in the messenger RNA molecule (mRNA) and the Iron Response Protein 1 (IRP1) [25–27]. In presence of high FIP level, IRP1 embeds an iron-sulfur cluster and adopts a closed conformation [27–29]. It has been demonstrated that the presence of the iron-sulfur cluster is fundamental to maintain the closed conformation of IRP1 [30–32]. Interestingly, the closed form of IRP1 represents the active form of the aconitase enzyme operative in the Krebs cycle [28,33,34], where it catalyzes the isomerization of citrate to *cis*-citrate via the *cis*-aconitate intermediate [35]. The lowering in FIP content shifts the chemical equilibrium, favoring the release of the embedded iron cluster. This, in turn, induces a conformational transition resulting in the fully opened and catalytically inactive form of IRP1. Importantly, the open conformation of IRP1 selectively interacts with the IRE stem-loop [36,37], leading to mRNA sequestration and, consequently, the downregulation of the ferritin production [38–41]. Indeed, IRP1-bound IRE is not able to recruit the small ribosomal unit, thus suppressing mRNA translation [25]. A reduction in ferritin levels also counteracts iron storage, hence leading to the increase of the FIP content until a break-even point is reached. Beyond this threshold, the iron-cluster is re-embedded in IRP1, resulting in the prevalence of the closed conformations (i.e. aconitase), the release of IRE, enhanced ferritin translation, and a subsequent decrease of FIP levels [2,42]. Recently, the free energy profile for the closed/open transition of IRP1 in presence or absence of an iron-sulfur cluster has been obtained using enhanced sampling molecular dynamic (MD) simulations confirming the crucial role of the ligand in shaping the free energy landscape [30]. Interestingly it has been shown that mutation-induced conformational changes in the IRE stem-loop altering the gene expression level [43].

**Fig. 1.**
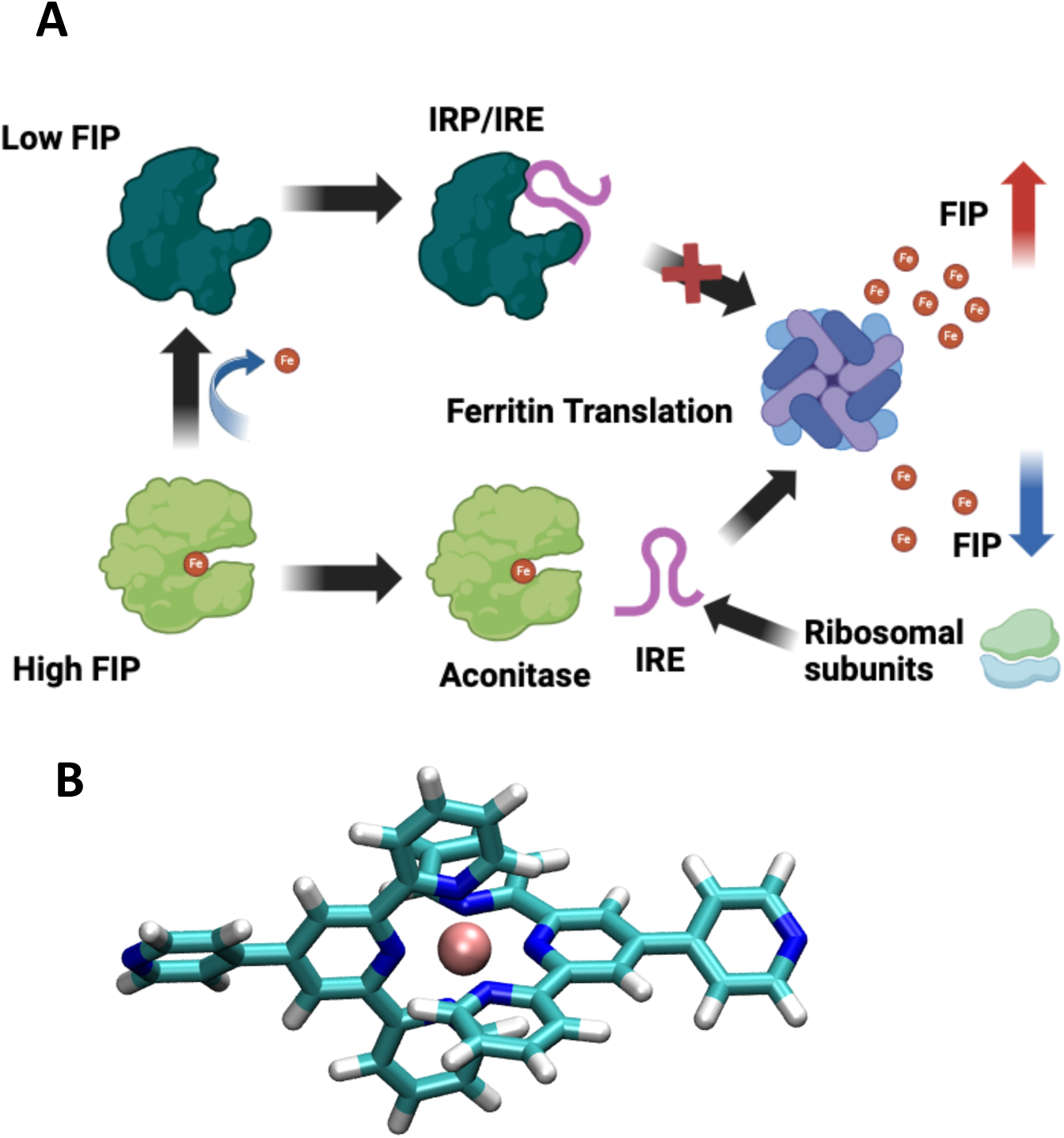
A) Schematic mechanism of the IRP-based regulation of iron hemostasis through the sequestration of the IRE mRNA. B) Structure of the AIM3 iron complex.

Iron complexes have shown increasingly appealing potential for biological and even clinical applications, particularly against cancer [44]. Some iron transition metals, bearing bipyridine units, in the following named AIM3 (Figure 1B), have been recently synthesized and studied aiming at understanding their unexpected properties and their biological effects. In particular, it has been shown that they can interact with double stranded DNA [45], are able to penetrate cellular membranes, reshape their morphology and mechanical properties [46], and may be related to the tuning of the epigenetic regulations. Furthermore, AIM3 has demonstrated antiproliferative capacities on cells [45]. These first observations questioned the binding behavior of the metal compound toward RNA non canonical motifs, mainly such as IRE stem-loop structures, and the biological consequences linked to a possible influence of the interaction between AIM3 and IRE on the regulation mechanism of the iron homeostasis. Interestingly it has been reported by Ke and Theil that a Copper phenantoline complex binds to IRE [47], suggesting conserved interactions patterns in different cell lines and thus its action as a iron-response protein regulator. For these reasons, we have performed UV/Vis, cellular biology assays, as well as long-range MD simulations, confirming that specific interactions of AIM3 with the IRE mRNA moiety can shape the regulatory response and alter the iron homeostasis cycle. However, we would like to stress out that our study, while it confirms the establishment of interactions between AIM3 and IRE, cannot discriminate on their specificity, whose validation would require additional experiments which clearly goes beyond the scope of the present contribution.

## 2. Material and methods

### UV-visible absorption

UV-vis spectra were recorded at room temperature on a Nanodrop 2000c spectrophotometer, from 190 to 800 nm. Lyophilized RNA were purchased from Eurogentec and resuspended in MilliQ water to yield 100 µM stock solutions. The IRE sequences of Ferritin H (5’ GGG GUU UCC UGC UUC AAC AGU GCU UGG ACG GAA CCC GGC GCU CGU 3’) and HIF2α (5’ UAC AAU CCU CGG CAG UGU CCU GAG ACU GUA 3’) were folded into IRE motifs by heating the solutions to 95°C for 5 min in 40 mM HEPES/KOH 100 mM KCl, pH 7.2, before slowly cooling to room temperature overnight. Titration experiments were carried out by adding increasing amounts of IRE RNA to a solution of AIM3 compound whose concentration was fixed to 100 µM.

### General cell culture

The human normal embryonic kidney 293T cell line (ATCC, Rockville, MD, USA) was cultured in DMEM (Sigma-Aldrich), with 10 % fetal bovine serum (Biowest), 2 mM L-glutamine (Sigma-Aldrich), and 0.1% Penicillin/Streptomycin antibiotics (Sigma-Aldrich). Cells were maintained in humidified atmosphere, at 37°C, with 5% CO_2_.

### Cell transfection

pYIC (Addgene plasmid #18673) and pIRE-YIC (Addgene plasmid #18784) plasmids were a gift from Han Htun [48]. 293T cells were seeded in a 6-well plate (3.0 x 10^5^ cells per well). Next day, culture medium was changed and cells transfected with 3 µg of plasmid DNA using jetPEI® DNA Transfection Reagent (Polyplus), following the manufacturer’s instructions. The day after, to evaluate a modulation effect on YFP translation, cells were treated with 50 or 100 µM deferasirox (DFX) (16753, Cayman Chemicals), 10 µM FeSO_4_ (215422, Sigma-Aldrich-Aldrich), or 2 or 10 µM AIM3 for 24 hours. YFP fluorescence was measured by analyzing living cells by flow cytometry using a CytoFLEX flow cytometer (Beckman Coulter, Life Sciences, B53000). Data were analyzed on the CytExpert software (Beckman Coulter). The percentage of YFP-positive cells was expressed relatively to the control condition, arbitrarily set to 1.

### Molecular dynamics simulations

To assess the interaction between IRP1/IRE and AIM3, we performed all-atom molecular dynamic simulations using NAMD code 3.0 [49,50]. The initial structure of the IRP1/IRE complex has been obtained from the X-ray crystallographic structure reported under the PDB code 3snp [37]. From this structure, different systems were prepared using AMBER software [51], and specifically the *tleap* module: the IRP1 protein surrounded by 15 AIM3 molecules; IRP1/IRE complex with one AIM3 unit placed in the protein groove between the protein and the mRNA; Apo IRE and IRE surrounded by 10 AIM3 molecules (see SI for the initial structures). All the systems were solvated in a cubic water box with 9 Å and described by using the TIP3P water model [52]. The Amber ff99bsc0 with χ-OL3 modification force field [53] was used to describe RNA, while Amber ff14SB on protein and the AIM3 force field was built based on generalized AMBER force field (gaff) as described in our previous work [45]. To ensure the electroneutrality of the systems, K^+^ cations were added. Moreover, to speed up the simulation, the H mass repartition (HRM) algorithm [54] was used in combination with the rattle and shake algorithms [55], allowing to scale up the time step from 2 to 4 ps. MD simulations were performed at a constant temperature (300 K) and pressure (1 atm) using the Langevin thermostat [56] and barometer [57], thus defining an isothermal and isobaric (NPT) thermodynamic ensemble. Particle mesh Ewald (PME) [58] was used to treat long range interactions with a cut-off of 9 Å. The following protocol was used for all the simulations: 5000 minimization steps to remove bad contacts, then the system was equilibrated during 9 ns with constraints applied in IRP1, IRE and AIM3 when present. During these first ns, the constraints were progressively decreased until none were applied. Afterward, the production runs were performed reaching 1.2 µs for Apo and IRE/AIM3 complex; 500 ns for the IRP1/AIM3 complex and 200 ns for the IRP1/IRE/AIM3 complex, with one replica for apo IRE, IRP1/AIM3 and IRP1/IRE/AIM3 complex, and three independent replicas for IRE/AIM3 complex.

All the analyses have been conducted using VMD [59] and *cpptraj* module of AMBER software, except for the RNA puckering analysis for which we used a home-made Python script as described in this work [60].

## 3. Results and Discussion

### Molecular Modeling and Simulations

The complex between IRE and IRP1 has been resolved by X-Ray crystallography, confirming the presence of a stable aggregate. The stability of this arrangement has also been confirmed by MD simulations reaching the μs timescale [30]. Interestingly, the proteins present two main contact points with the mRNA strand involving either the loop of the messenger RNA or the unpaired nucleobases, mainly uracil. In contrast, the central double helical region remains rather solvent exposed. Interestingly, such interaction pattern may also offer enhanced sequence and structural selectivity in agreement with the IRP1 biological role. In a previous work, using MD simulation, we have also confirmed the main binding sites of IRP1 with IRE, which mainly involve nucleotides in the loop or the edge region of the mRNA [30]. Interestingly, in the closed aconitase-like conformation the IRE-binding region is not exposed precluding the interaction with the IRE RNA stem-loop.

To rationalize the effects of the inclusion of AIM3, we have also modeled the interaction of the transition metal complex with IRP1 and with the already formed IRP1/IRE complex. As reported in SI (Figure S1), we have shown that AIM3 may bind to the surface of IRP1, as expected due to the coexistence of a net charge and hydrophobic ligands. Yet, no specific interaction may be evidenced in the two main recognition sites of IRE. The same behavior can be observed in the case of the simulations for the pre-formed IRP1/IRE aggregate in presence of AIM3 (Figure S2). Indeed, while some specific interactions of AIM3 are evidenced, they take place in a region of the complex which is unlikely to induce structural deformation, and thus perturb the protein/RNA binding.

The situation is instead totally different in case of the IRE mRNA fragment. Indeed, in three independent replicas, we have shown that AIM3 is able to perform stable and rather specific interactions with different RNA spots as shown in Figure 2. Importantly, we should stress out that the AIM3/IRE aggregates are formed entirely spontaneously in a very short time span and that they lead to stable aggregates persisting for the whole range of the MD simulation, i.e., above the μs timescale. From Figure 2 A and B, it appears clearly that on the one side the interaction with IRE is not inducing significant and strong structural deformation of the mRNA strand, and most notably is conserving the stem-loop arrangement. However, it is also evident that IRE can interact with up to five AIM3 molecules which occupy different interaction sites. Unsurprisingly, and due to the positive charge of AIM3, the ligand as a tendency to slide into the mRNA groove, where it is stabilized with favorable interactions with the negatively charged phosphate of the backbone, as previously observed with DNA double strand [45]. Yet additional π-stacking between the uncoupled mRNA bases and the conjugated AIM3 ligands also takes place further stabilizing the complex. Importantly, we may recognize that the transition metal ligand is favorably placed in the loop region marked as region 1 in Figure 2B as well as at region encompassing the extruded bases (2 in Figure 2B) and at the stem-loop hedge (3 in Figure 2B).

**Fig. 2.**
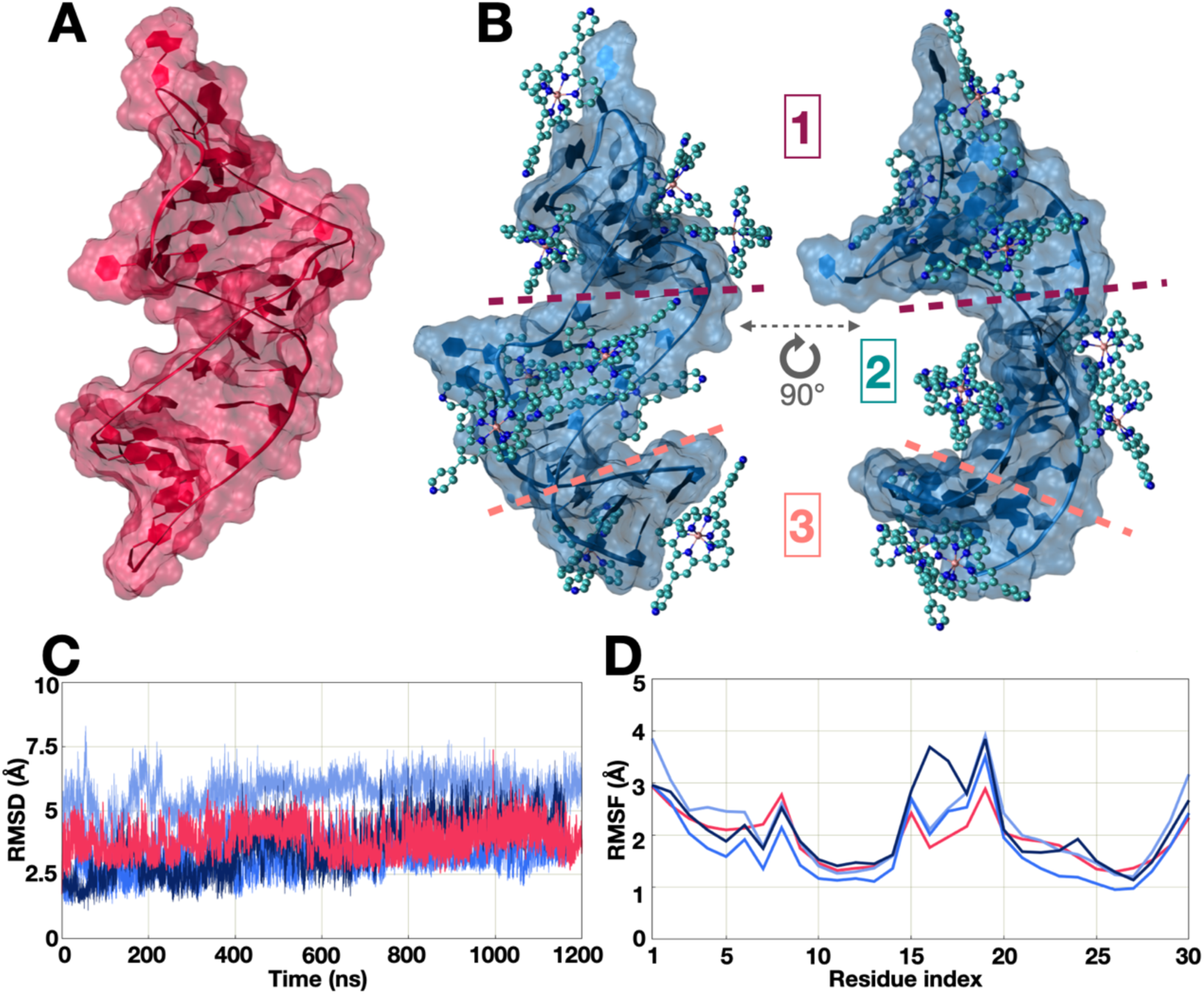
A representative snapshot extracted from the MD simulation for the APO IRE (A) and the IRE/AIM3 complex (B). RMSD (C) and RMSF (D) for the APO mRNA (in red) and for the three replicas for the complex (in blue).

The structural stability of the complex and the absence of pronounced structural deformations can also be appreciated from the analysis of the RMSD time-series for the IRE/AIM3 complex which is almost perfectly overlapping with the one of the APO IRE (Figure 3 C). The RMSF for the mRNA (Figure 3 D) also shows that the interaction with AIM3 is not significantly altering the flexibility profile since the variations in the residue fluctuations does not exceed 1 Å compared to the APO system. Yet, the larger variations seem to involve the mRNA loop region, i.e., one of the most favorable binding spot see figures.

**Fig 3.**
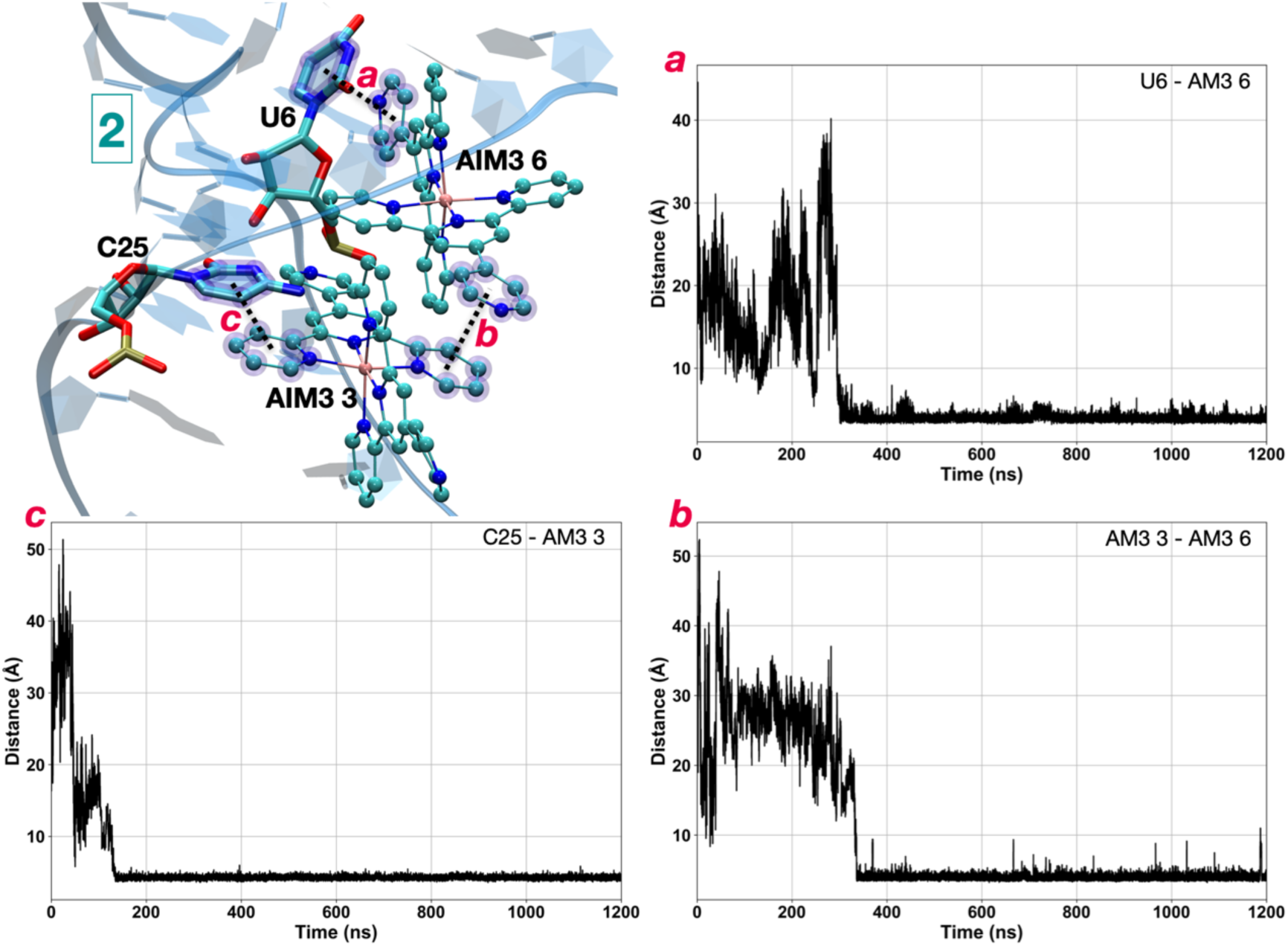
A representative snapshot of AIM3 interacting in the extruded nucleobases region of IRE. Time series of the distances between AIM3-5 and U6 (a), AIM3-6 and C25 (b) and between the two AIM3 ligands (c).

In Figure 3 we report a representative snapshot and the timeseries of the most crucial distances for AIM3 interacting in the extruded nucleobases region. We may clearly evidence that strong interactions takes place between AIM3 and either C25 or U6 establishing in the 200-400 ns range and persisting all along the MD simulation. Interestingly these interactions give raise to a stable arrangement in which two AIM3 ligands occupy a proximal region and develop favorable π-stacking between their aromatic ligands. potentially shielding the RNA extruded nucleobases from the interaction with IRP1. Similar interactions also take place on the other regions of the stem-loop as reported in Figures S4-S6. These findings are also coherent with the results reported in reference [47], which prove the interaction of a Cu complex bearing extended π-conjugated ligands with IRE. Although, no molecular and atomic-scale resolution structural determination of the transition metal complex with IRE was provided in [47].

However, the global stability as observed from the RMSD may mask more subtle structural variations, induced by the interaction with AIM3 and mostly related to the ribose puckering. The former parameter is a key RNA structural indicator, which is also strongly related to the nucleic acid intrinsic reactivity and to its capacity to bind different macromolecular partners. As reported in Figure 4, we can clearly see that the conformational equilibrium, especially for the loop regions (nucleotides 15 to 19) is strongly altered. In all the three replicas, we may observe that the C2’-*endo* (B-like family) conformation of the sugar is strongly reduced upon interaction with AIM for C18 and to a lesser extent for U17. On the other hand, U19 increases the presence of the C2’-*endo* form. As already pointed out, these residues are extruded nucleobases which are forming the loop region and are strongly involved in the interaction with AIM3. Interestingly, the same puckering modification can be observed for the other extruded nucleobases, such as G7, for which the C2’-*endo* conformation is dominant in the APO form but almost disappears upon interaction with AIM3 at the profit of the C3’-*endo* form (A-like family). Interestingly, this modification induces an increase of the C2’-*endo* conformation for the nearby residues U6 and C8.

**Fig. 4.**
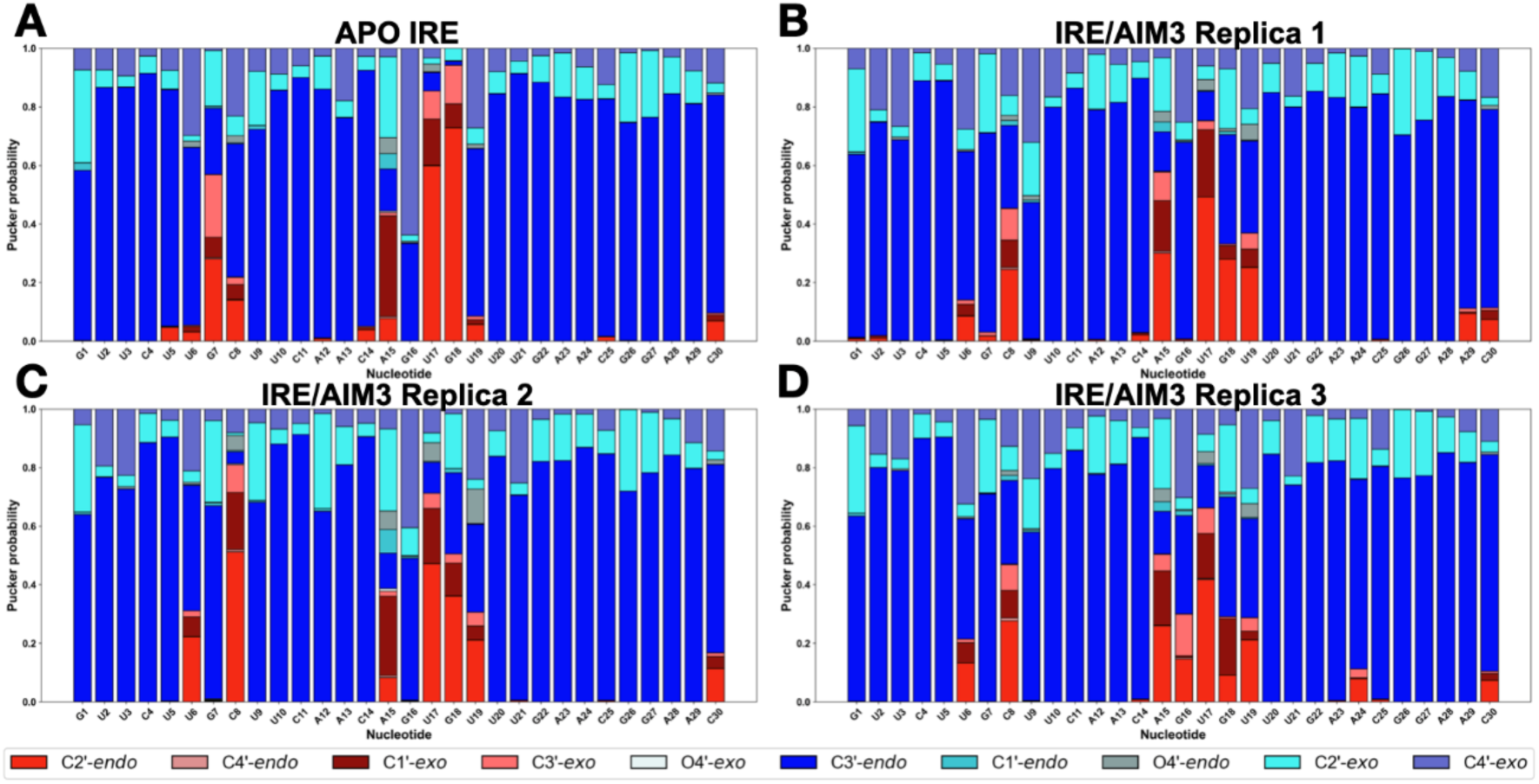
Stacked bar plot representing the pucker probability of apo IRE (A) and the three independent replicas of the IRE/AIM3 complex (B, C, D). B-like family: Red: C2’-*endo*, light pink: C4’-*endo*, dark red: C1’-*exo*, light red: C3’-*exo*. A-like family: Blue: C3’-*endo*, dark cyan: C1’-*endo*, cyan: C2’-*exo*, light blue: C4’-*exo*, grey: O4’-*endo*, white: O4’-*exo*.

### UV/Vis

To experimentally confirm the capacity of AIM3 to interact with different IRE mRNA motifs in solution, we performed UV-visible titrations. The characteristic absorption bands in the UV-vis spectrum of AIM3 in the regions which are not overlapping with RNA absorption (324 nm and 570 nm) were monitored in presence of increasing quantities of IRE motifs. The 5’ UTR regions of *Ferritin H* [10] and *HIF2α* [61] mRNAs were considered for analysis since they represent well-characterized IRE structures.

The increase of the concentration of the mRNA titrants leads to significant modifications of the AIM3 spectrum, primarily resulting in a hypochromic effect, as presented in Figure 5. Specifically, both characteristic bands of AIM3 exhibited approximately 18% and 25% hypochromism upon addition of *Ferritin H* and *HIF2α* mRNA sequences, respectively. However, a distinct trend was observed between the two stem-loop mRNA fragments: the addition of *HIF2α* IRE sequence induced a progressive, dose-dependent decrease of AIM3 absorbance, whereas the addition of the *Ferritin H* oligonucleotide caused an immediate hypochromic effect upon its first addition, without further dose-dependent response. Additionally, increasing concentrations of *Ferritin H* provoked a slight red shift of the 570 nm absorption band in the AIM3 spectrum (Figure 5 A). Overall, the different spectral changes observed upon the incremental addition of RNA oligonucleotides demonstrate the capacity of AIM3 to interact with IRE mRNA motifs in solution, confirming the results obtained by MD simulations.

**Fig. 5.**
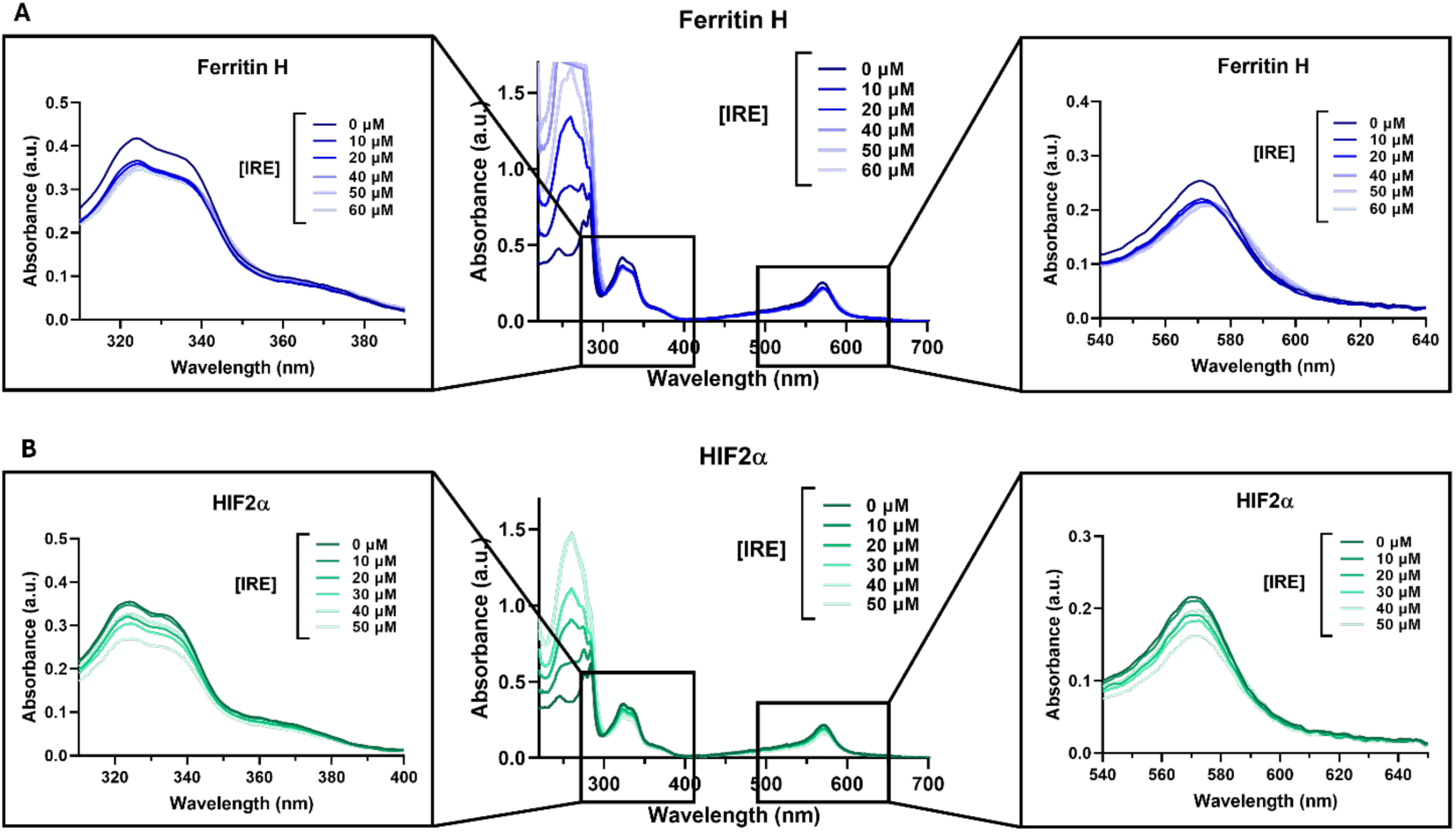
UV-vis spectra of AIM3 in presence of increasing amounts of RNA. The IRE motif of Ferritin H (A), and HIF2α (B) were considered to titrate AIM3 (100 µM), in 40 mM HEPES/KOH, 100 mM KCl, pH 7.2. [IRE] = 0 – 60 µM.

### Cellular Assays

In the previous sections, we have demonstrated that AIM3 interacts with the IRE motif, possibly leading to its stabilization. Such stabilization may enhance its accessibility to IRP1 partners, thereby promoting transcript repression. Thus, to further characterize the effects of AIM3 – IRE interactions in a cellular environment, cells have been transiently transfected with an IRE-containing gene reporter system, namely the pIRE-YIC plasmid. This plasmid encodes the yellow fluorescent protein (YFP), whose expression is under the control of a 5’ UTR mRNA sequence harboring an IRE motif. Consequently, YFP expression directly correlates with the intracellular free iron pool (FIP), serving as a witness of translation regulation by the IRE/IRP1 system.

In this respect, in presence of high FIP, IRP1 proteins sequester iron and are converted into aconitase form, which is unable to bind IRE motifs on the 5’ UTR mRNA, thereby allowing YFP expression. In contrast, when FIP is low, IRP1 act as repressors of translation leading to reduced YFP expression.

As a control, the pYIC plasmid was used, as its 5’ UTR sequence does not fold into an IRE structure. Therefore, YFP translation occurs independently of the IRE cap-dependent repression. A mock condition, in which plasmid DNA was replaced with water, was also included as an additional control.

As a further control, cells were also treated with the iron chelator deferasirox (DFX), which is expected to reduce FIP levels, promote IRP1 binding to IRE, and consequently reduce YFP expression. In cells transfected with the IRE-containing plasmid and treated with DFX, the proportion of YFP-positive cells decreased by approximately 1.2-fold compared to the control; however, this reduction did not reach statistical significance. Conversely, the addition of iron via FeSO_4_ treatment significantly increased the proportion of YFP-positive cells, with a fold change of 1.4 (Figure 5). These results confirm the sensibility of the pIRE-YIC plasmid to the IRP1/IRE-based translational regulation.

Interestingly, AIM3 treatment led to a significant reduction in the proportion of YFP-positive cells, which decrease by approximately 1.5-fold at 2 µM and 1.8 at 10 µM. In contrast, in the absence of an IRE motif in the 5’ UTR sequence (pYIC plasmid), no statistically significant changes were observed in the YFP expression (Figure 6). Therefore, our results indicate that AIM3 functions as a repressor of translation only in presence of folded IRE motif in the 5’ UTR mRNA region. The differential effect of AIM3 depending on the presence of IRE motif hints its direct involvement, likely through its previously demonstrated capacity to bind to the 5’ UTR mRNA region.

**Fig. 6.**
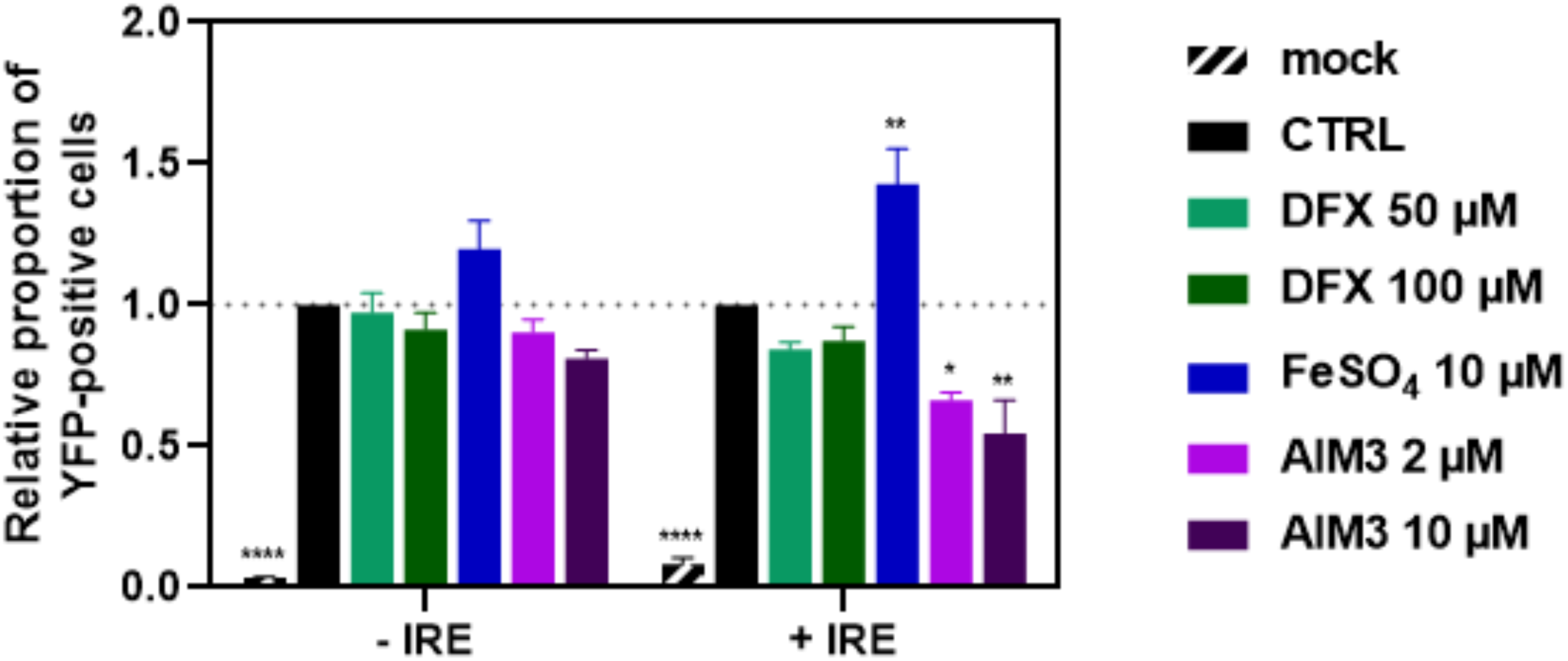
Effects of AIM3 treatment on the translation of an IRE-containing mRNA. 293T cells were transiently transfected with a no IRE-containing reporter gene (-IRE, pYIC DNA plasmid), or an IRE-containing reporter gene (+ IRE, pIRE-YIC DNA plasmid), and treated with 50 or 100 µM deferasirox (DFX), 10 µM FeSO_4_, and 2 or 10 µM AIM3 compound. In mock condition, plasmid DNA was replaced by water. YFP expression was evaluated by flow cytometry. Mean ± SEM (n=3). One-way ANOVA, Dunnett’s multiple comparison test.

The formation of stable IRE/AIM3 aggregates are therefore coherent with a stabilization of the RNA structural motif favoring its binding with IRP1. This hypothesis is further supported by AIM3 ability to interact with the pre-formed IRP1-IRE complex without disrupting its stability, as confirmed by MD simulations. Indeed, a recent report describes that the small molecule Posiphen acts as a downregulator of the APP and HTT proteins by binding to the already formed IRP1 – IRE complex [62]. Furthermore; 5’ UTR regions are key regulatory sequences in RNA that play a crucial role in translation initiation, mainly by facilitating the recruitment of the 40S pre-initiation complex. IRE structures within these regions have been shown to be essential for this process [25,63]. Our findings indicate that AIM3 primarily binds to the stem loop of the IRE but can also aggregate at multiple interaction sites, with up to five molecules of the transition metal compound binding to the IRE. This suggests that, in addition to promoting IRP1 recruitment to the IRE, AIM3 may also hinder the recruitment of the 40S ribosomal subunit because of steric hindrance, thereby inhibiting translation initiation.

## 4. Conclusions

The regulation of iron homeostasis is a complex process which is dynamically regulated at the translational level via the IRP1/aconitase equilibrium and the correlated sequestration of the ferritin mRNA via specific interactions with its IRE stem-loop We have shown, by using a combination of molecular modeling and simulation and experimental techniques, including UV/vis titration, that AIM3 transition metal complex is able to bind to the IRE messenger RNA stem loops. The formation of the aggregate is mainly driven by electrostatic interactions with the nucleic acid negative groove and by the formation of favorable π-stacking interactions also involving extruded nucleobases. Importantly, we have shown that the binding does not induce pronounced structural deformations of the nucleic acid conformations. Additionally, we have also shown that the presence of AIM3 is not disrupting a preformed IRP1/IRE complex, coherently with the small structural perturbations induced by the ligands. Finally, we have shown that in cells treated with AIM3, the translation of IRE-dependent proteins is strongly downregulated, with a dose-dependent effect which is even higher than the one induced by iron chelators. Although further studies will be necessary to clearly elucidate this mechanism, we can hypothesize that the downregulation could be due either to the stabilization of the IRE motif which favors its binding to and sequestration by IRP1, and/or to the inhibition of the downstream recruitment of ribosomal factors due to the presence of AIM3. Our results unambiguously show that AIM3 is an efficient regulator of iron hemostasis. Interestingly, this regulation is not induced by direct iron chelation, but rather through the interaction with regulatory mRNA elements, influencing the ferritin translational regulation.

Recently, the scientific community has emphasized the possible development of novel small molecules to target proteins that are considered difficult to drug due to the lack of well-defined druggable pockets. The main emerging strategy to overcome this challenge is the direct repression of protein translation by targeting the corresponding mRNA. Thus, small molecules have been designed to selectively bind to particular tridimensional structures found in transcripts, including IRE motifs [64]. Although only a few such molecules have been reported to specifically target IRE yet, some studies report the ability of such small IRE ligands to regulate protein translation, either positively or negatively [43,61,62,65,66]. It is important to underline, however, that our results do not indicate a specific and selective interaction between IRE and AIM3, in particular concerning possible mutations of IRE which may alter its recognition [47]. To answer this question additional experiments, involving in particular control sequences should be performed at the experimental and MD level, which could be the object of forthcoming contributions.

Thus, our study may represent an important milestone in the development of specific RNA-targeting drugs for the control of iron hemostasis. In the future, we plan to further characterize the binding free energy between IRE and IRP1 in presence of AIM3. Furthermore, we also plan to study the downstream interaction of IRE with ribosomal subunits to assess the potential inhibition of this interaction by AIM3.

## Acknowledgments

The authors thank GENCI and Explor computing centers and the Platform P3MB for computational resources. The authors thanks ANR and CGI for their financial support of this work through Labex SEAM ANR 11 LABEX 086, ANR 11 IDEX 05 02 and PIRATE. The support of the IdEx “Université Paris 2019” ANR-18-IDEX-0001.

## Electronic supplementary materials

Analysis of the IRP/AIM3 and of the IRP/IRE/AIM3 complexes. Detailed analysis of the interactions between AIM3 and the IRE extruded nucleobases. (PDF file).

## Supporting Information

**Fig. S1.**
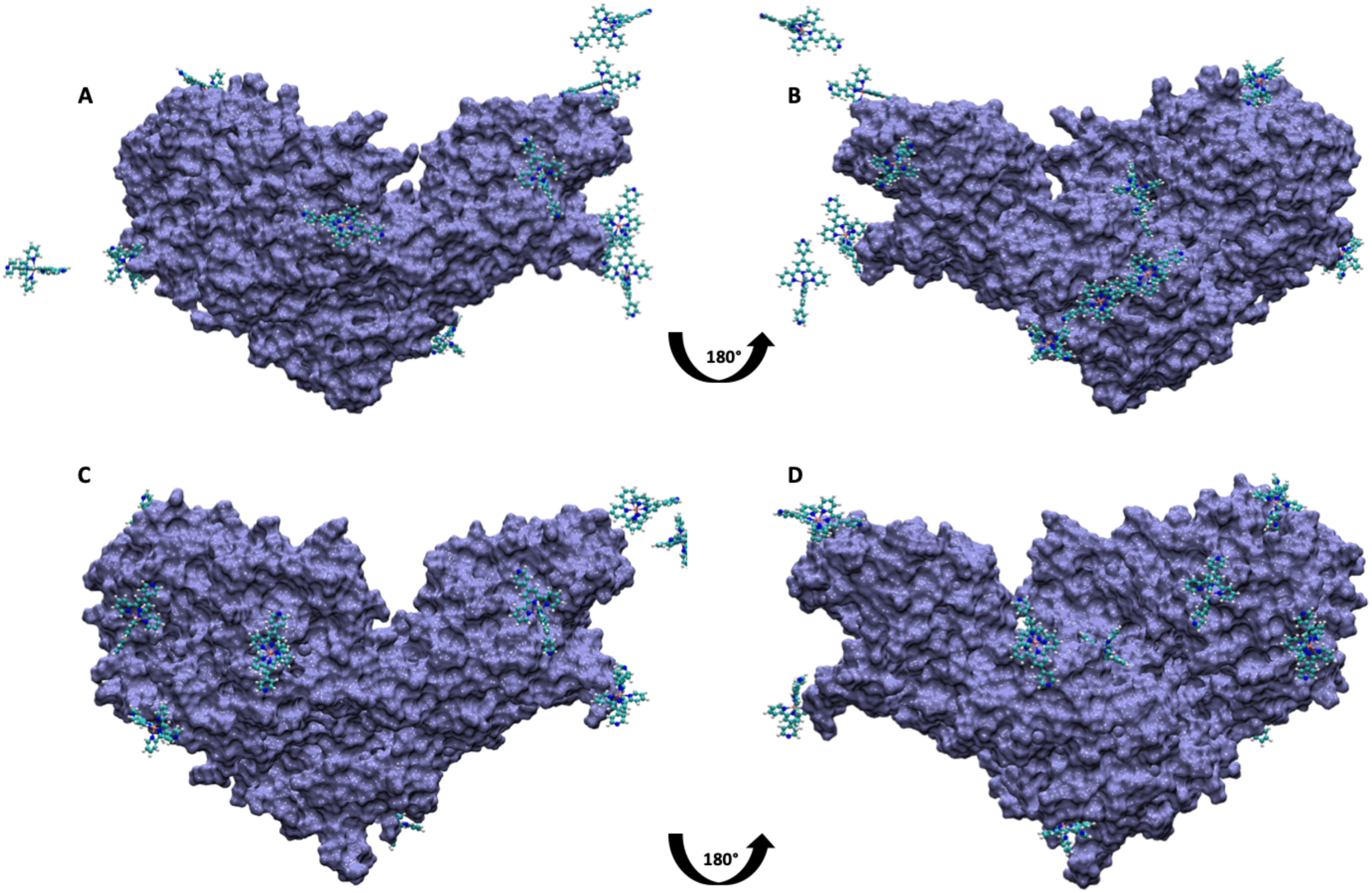
Interaction between IRP and AIM3 at 245 ns (A and B) and at 500 ns (C and D). A(B) and C(D) represent the front and back view of IRP respectively.

**Fig. S2.**
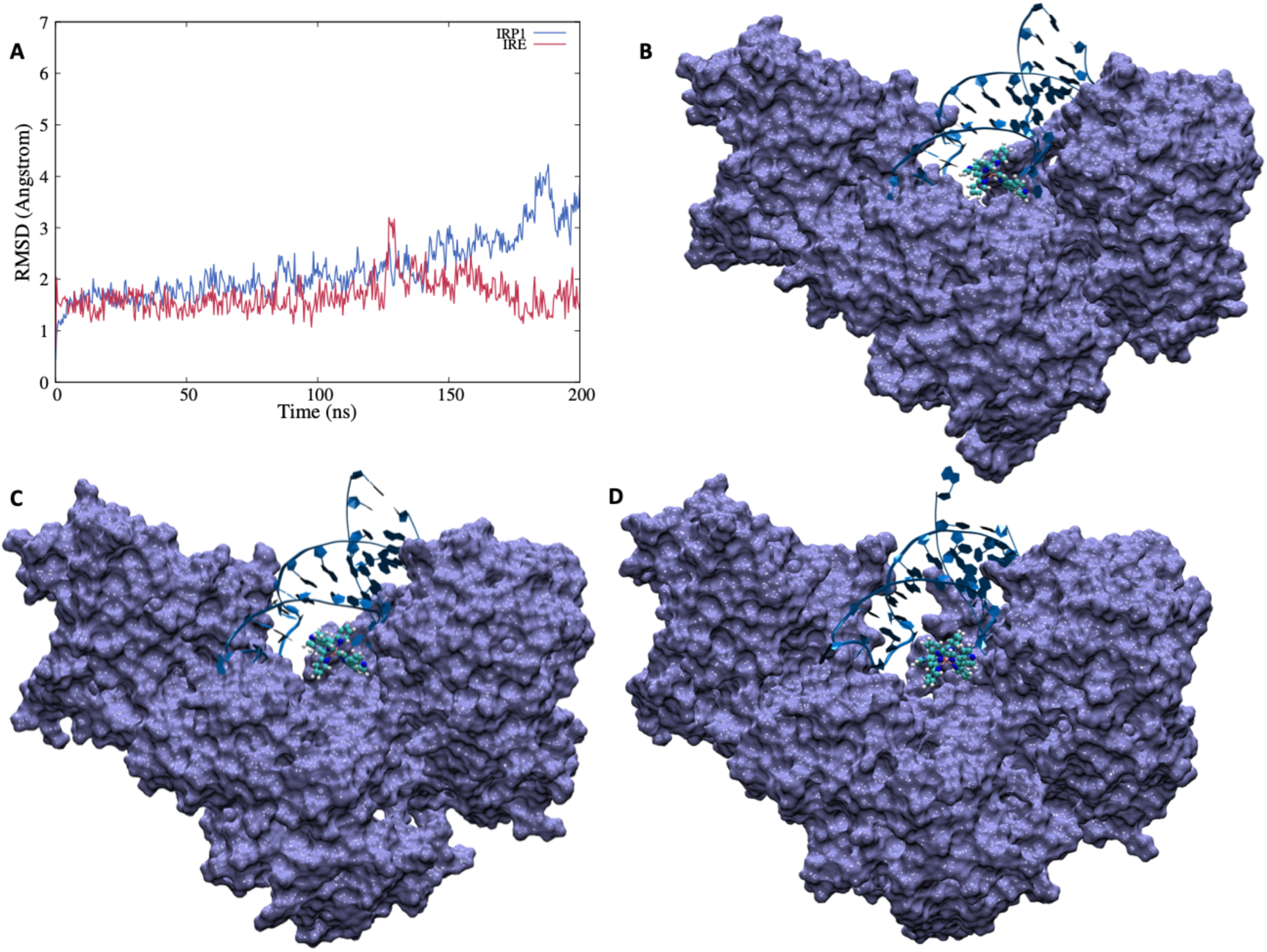
Interaction between the preformed IRP/IRE complex and AIM3 A) time series of the protein and RNA RMSD. Time evolution of the structure at 0 ns (B), 100 ns (C) and 200 ns (D).

**Figure S3.**
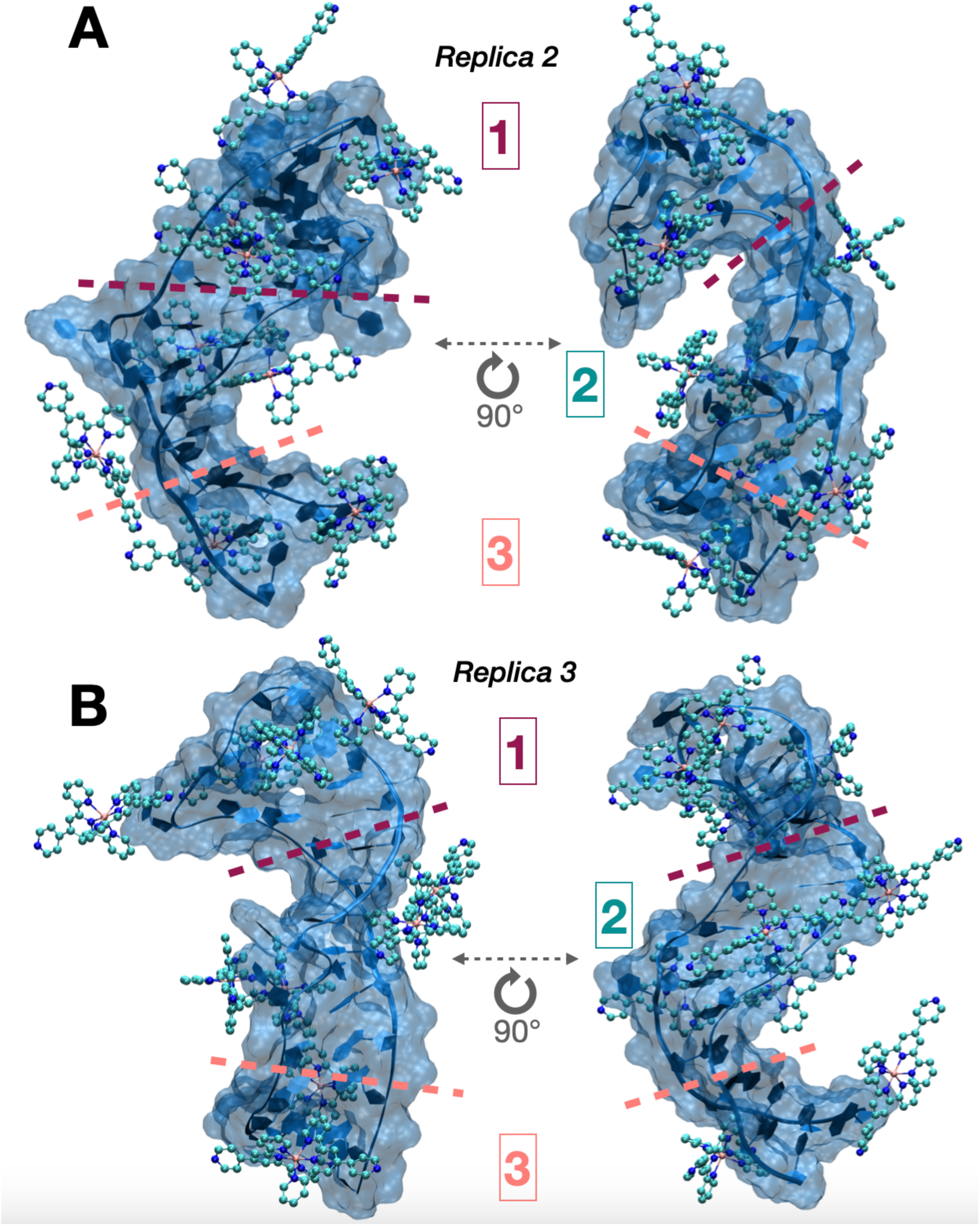
Interaction between AIM3 and IRE as obtained for Replica 2 and Replica 3.

**Fig. S4.**
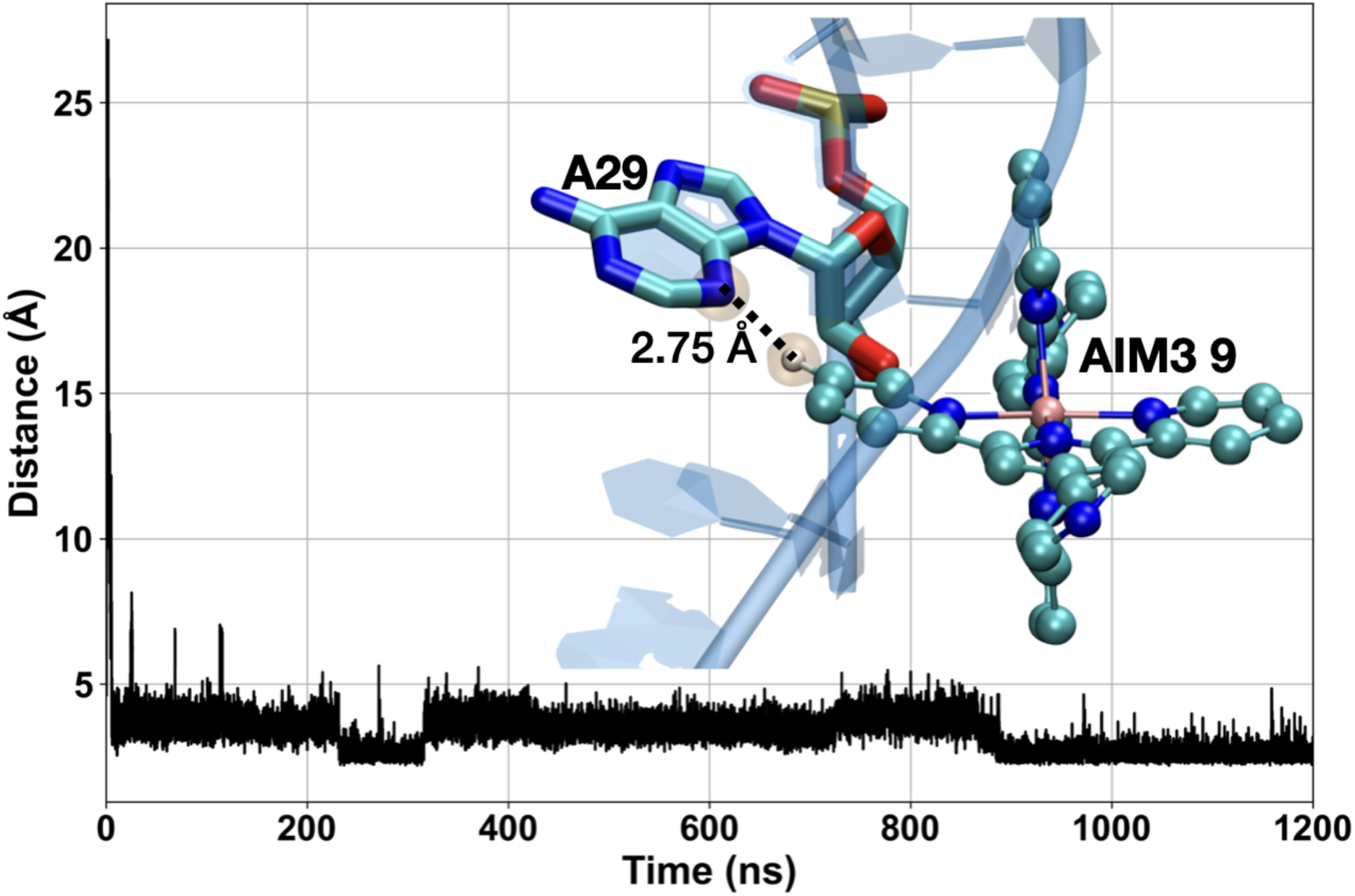
Time evolution of the distance between H19 of AIM3 9 and N3 of nucleotide A29 with a representative snapshot of the interaction. Atoms implied in the bond are highlighted in orange.

**Fig. S5.**
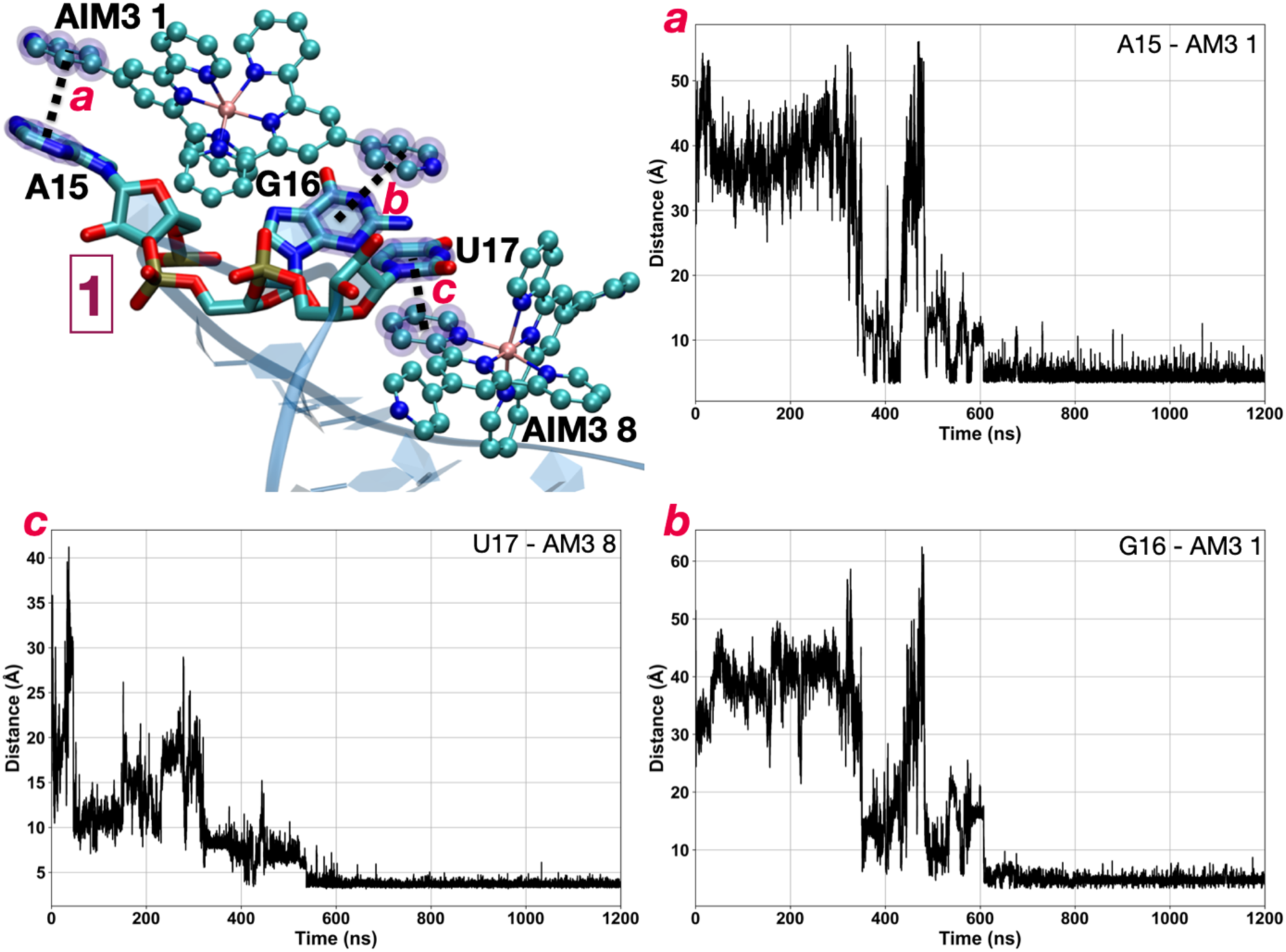
Time evolution of the distance between AIM3 1 and AIM3 8 and nucleotides A15, G16 and U17 a) distance between A15 and AIM3-1, b) distance between G16 and AIM3-1, c) distance between U17 and AIM3-8, and.

**Fig. S6.**
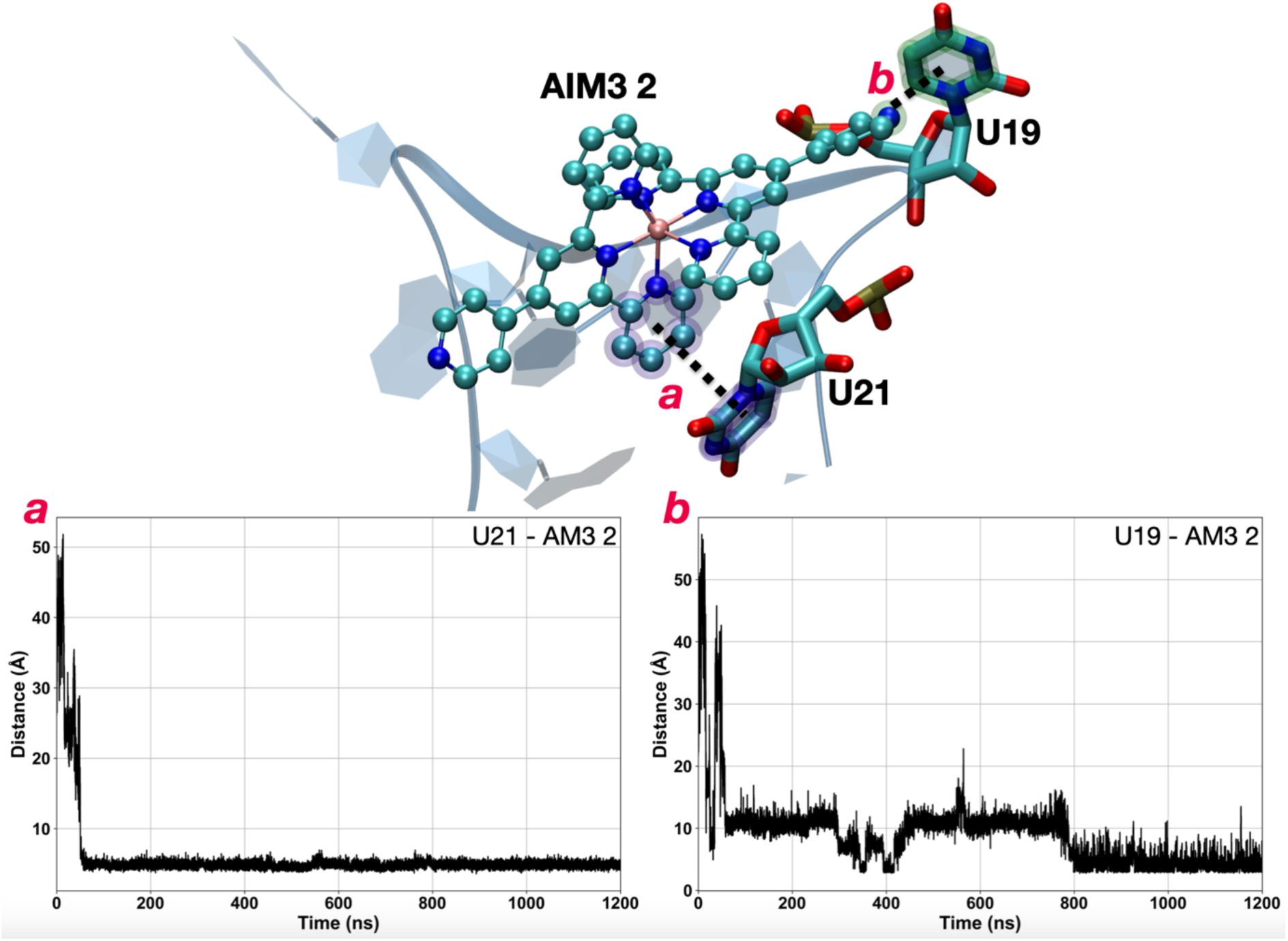
Time evolution of the distance between AIM3 2 and nucleotide U21 or U19 with a representative snapshot of the interactions. Atoms implied in the bond are highlighted in green and violet.

**Fig. S7.**
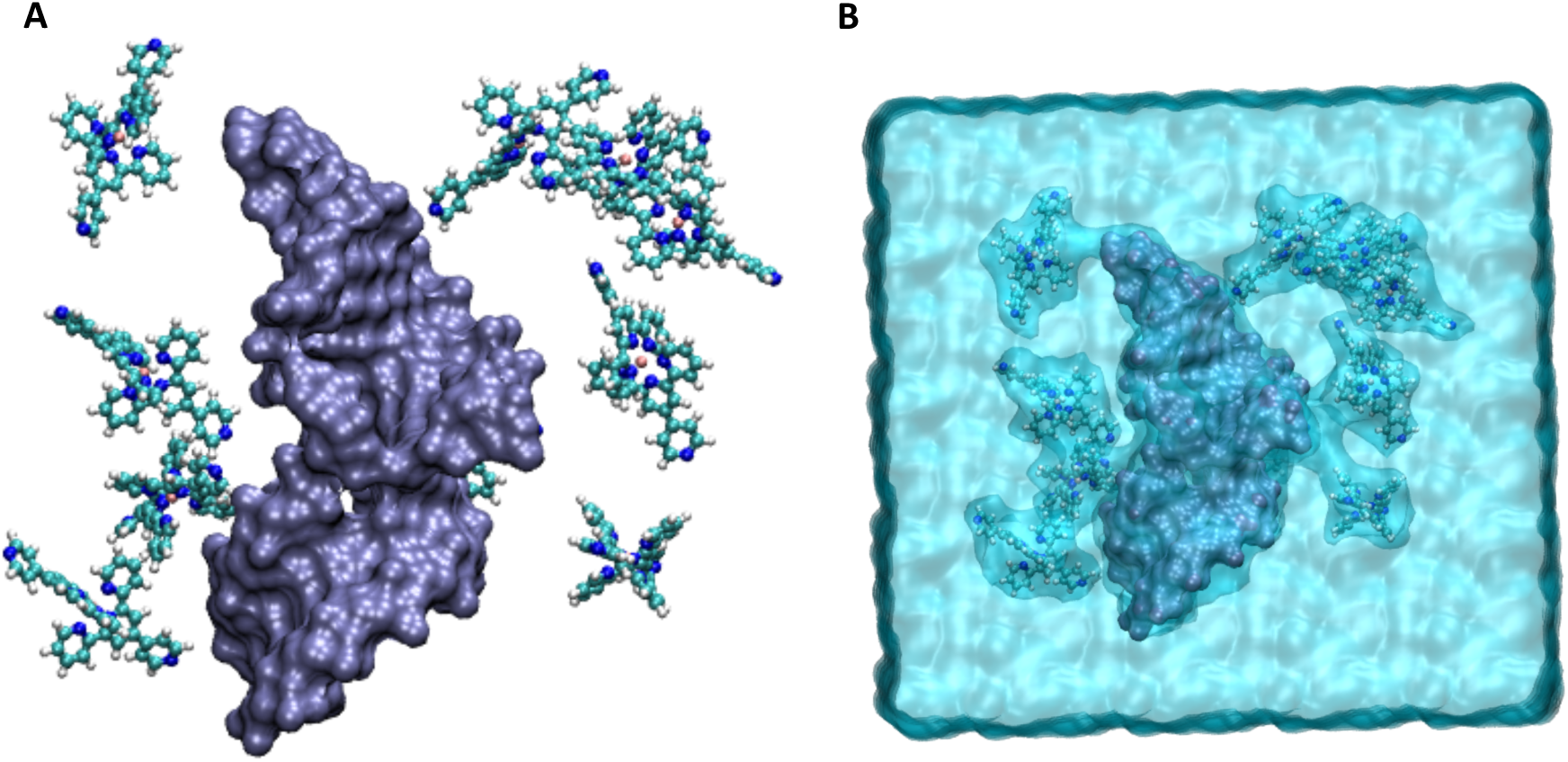
A) Initial structure of the IRE stem-loop (surface) interacting with 10 AIM3 balls and sticks). B) The initial system including.

